# Analysis of Normal Levels of Urine and Plasma Free Glycosaminoglycans in Adults

**DOI:** 10.1101/2021.05.21.445098

**Authors:** Sinisa Bratulic, Angelo Limeta, Francesca Maccari, Fabio Galeotti, Nicola Volpi, Max Levin, Jens Nielsen, Francesco Gatto

## Abstract

Plasma and urine glycosaminoglycans (GAGs) are long linear sulfated polysaccharides recognized as potential non-invasive biomarkers for several diseases. However, owing to the analytical complexity associated with the measurement of GAG concentration and disaccharide composition, the so-called GAGome, a reference study of the normal healthy GAGome is currently missing. Here, we prospectively enrolled 308 healthy adults and analyzed their urine and plasma free GAGomes using a standardized ultra-high-performance liquid chromatography coupled with triple-quadrupole tandem mass spectrometry (UHPLC-MS/MS) method together with comprehensive demographic and blood chemistry biomarker data. Of 25 blood chemistry biomarkers, we mainly observed weak correlations between the free GAGome and creatinine in urine, and hemoglobin or erythrocyte counts in plasma. We found higher free GAGome concentration – but not composition - in males. Partitioned by gender, we established reference intervals for all detectable free GAGome features in urine and plasma. We carried out a transference analysis in healthy individuals from two distinct geographical sites, including the Lifelines Cohort Study, which validated the reference intervals in urine. Our study is the first large-scale determination of normal plasma and urine free GAGomes reference intervals and represents a critical resource for physiology and biomarker research.

## Introduction

Glycosaminoglycans (GAGs) are a family of long linear polysaccharides consisting of repeating disaccharide units (*1*). Different classes of GAGs have been characterized. In humans, the most prevalent classes are chondroitine sulfate (CS) [(→3)-β-D-GalNAc(1→4)-β-D-GlcA or α-L-IdoA(1→], heparan sulfate (HS) [(→4)-α-D-GlcNAc or α-D-GlcNS(→4)-β-D-GlcA or α-L-IdoA (1→], and hyaluronic acid (HA) [(→3)-β-D-GlcNAc(1 → 4)-β-D-GlcA(1→] where GalNAc is N-acetylgalactosamine, GlcA is glucuronic acid, IdoA is iduronic acid, GlcNAc is N-acetylglucosamine, and GlcNS is N-sulfoglucosamine. glucuronic acid, which can be further modified by sulfation in up to three sites. CS and HS disaccharides can each be further modified with O-sulfo groups in up to three positions. The resulting sulfation motifs confer GAGs highly diverse biological functions that are essential for healthy human development and physiology (*2*). The panel of GAG motifs resulting from the diversity in structure and concentration of GAGs is collectively referred to as GAGome.

Alterations in the physiological function of GAGs have been associated with several diseases ranging from mucopolysaccharidosis, a group of rare metabolic disorders caused by genetic defects in lysosomal enzymes that degrade GAGs, to complex diseases such as sepsis, rheumatoid arthrosis, and cancer (*3–6*). Plasma and urine GAGomes have been proposed as promising biomarkers for early non-invasive diagnostics (*6*). Despite the potential role of GAGs for clinical applications, the measurements of the GAGome has been limited to very small sample sizes (ranging from 3 healthy donors in (*7*) to 25 in (*6*)), in predominantly retrospective and selected donors, with different analytical techniques performed within academic laboratories (*3*, *4*, *6–9*). These limitations are due to the historical lack of effective analytical methods until recently (*10–14*), which proved hard to standardize and expensive to run. As a result, the GAGome values reported in the literature for healthy subjects are widely variable and cannot be consistently used as reference for physiology and biomarker research.

In this study, we took advantage of a standardized analytical method using ultra-high-performance liquid chromatography coupled with triple-quadrupole tandem mass spectrometry (UHPLC-MS/MS) (*15*) to analyze the urine and plasma free GAGomes in a Good Laboratory Practice (GLP)-compliant blinded central laboratory in two prospectively enrolled independent cohorts of 308 self-reported healthy subjects with comprehensive demographic and blood chemistry data. We first determined the correlation of free GAGomes with demographic and blood chemistry variables. Next, we established reference intervals for the normal urine and plasma free GAGomes according to accepted guidelines (CLSI EP28-A3c). Finally, we validated the proposed references intervals by transference analysis on two independent cohorts consisting of a total of 140 healthy individuals from two distinct geographical sites.

## Patients and Methods

### Study design

This study was designed and conducted in compliance with the CLSI. Defining, Establishing, and Verifying Reference Intervals in the Clinical Laboratory; Approved Guideline—Third Edition. CLSI document EP28-A3c. Wayne, PA: Clinical and Laboratory Standards Institute, 2008. The collection of specimens was planned prospectively with *a priori* criteria for population sampling. The present study was approved by the Ethical Committee (Etikprövningsmyndigheten) in Gothenburg, Sweden (#737-17 and #198-16).

### Reference sample group population

This study prospectively enrolled self-rated healthy adult subjects in one site in Sweden forming two independent cohorts. Cohort 1 and 2 were used to form the reference sample group and they were both enrolled at Sabbatsberg Hospital, Stockholm, Sweden between May 2018 and December 2019. Inclusion criteria were: adults between 21 and 78 years old; at least moderate self-rated health; no history of cancer (except non-malignant skin cancer); no family history of cancer (first-degree relative); fit to undergo protocol procedures. Exclusion criteria were: abnormal PSA value in the last 5 years. Eligible participants were identified among volunteers based on a questionnaire by trained research nurses. Participants in each cohort formed a consecutive series. EDTA-plasma and spot-urine samples were collected in one single visit. Blood samples were also obtained by venipuncture from participants in Cohort 1 and 2 to evaluate laboratory biomarkers informative of the general health status of the subject, including the complete blood count, and the concentration of electrolytes (sodium, potassium, calcium), ALAT, ASAT, CRP, PSA, glycated hemoglobin (Cohort 2 only), LDL and HDL (Cohort 1 only). Subjects with abnormal values were referred for clinical examination but were otherwise retained in the study.

### Transference population

The transference analysis was performed on two cohorts (Cohorts 3 and 4) representative of healthy adults from two distinct and external geographical sites. Cohort 3 used retrospectively archived specimens from the Lifelines Cohort study (*16*). Inclusion criteria were: adults older than 18 years old; self-reported healthy. Exclusion criteria were: diagnosis of cancer within 18 months from sampling visit. Lifelines is a multi-disciplinary prospective population-based cohort study examining in a unique three-generation design the health and health-related behaviours of 167,729 persons living in the North of the Netherlands. It employs a broad range of investigative procedures in assessing the biomedical, socio-demographic, behavioural, physical and psychological factors which contribute to the health and disease of the general population, with a special focus on multi-morbidity and complex genetics. Cohort 4 was enrolled prospectively at Sahlgrenska University Hospital, Göteborg, Sweden. Inclusion criteria were: adults older than 18 years old; self-reported healthy. Exclusion criteria were: none.

### Glycosaminoglycan measurements

Urine was collected at room temperature in a polypropylene cup. EDTA-plasma was collected through venipuncture in a vacuette at room temperature and next subjected to centrifugation (1100-1300 g, 20 minutes at room temperature in Cohort 1, 2500 g 15 minutes at 4 °C in Cohort 2) within 15 minutes. Samples could be stored refrigerated (4 °C) for 12 hours prior to transfer to a freezer (−20 °C in Cohort 1, −70 °C in Cohort 2). Shipment was performed at the same temperature as storage.

Sample preparation was performed in a single blinded GLP-compliant laboratory using the MIRAM™ Free Glycosaminoglycan Kit (Elypta AB, Sweden) to extract glycosaminoglycans (GAGs) from urine samples. GAG detection and quantification was obtained through ultra-high-performance liquid chromatography coupled with triple-quadrupole mass spectrometry (Waters® Acquity I-class Plus Xevo TQ-S micro) in accordance with the instruction for use in of the kit (*15*).

Laboratory measurements of the GAGome included the absolute concentration in μg/mL of chondroitin sulfate (CS), heparan sulfate (HS), hyaluronic acid (HA) disaccharides, resulting in 17 independently measured features. Specifically, we quantified eight CS disaccharides (0s CS, 2s CS, 6s CS, 4s CS, 2s6s CS, 2s4s CS, 4s6s CS, Tris CS) and eight HS disaccharides (0s HS, 2s HS, 6s HS, Ns HS, Ns6s HS, Ns2s HS, 2s6s HS, Tris HS) – corresponding to different sulfation patterns of CS and HS – and one HA disaccharide. A measured GAGome feature was considered detectable in a fluid (plasma or urine) if its mean concentration across all samples was > 0.1 ug mL^-1^(*15*).

The detectable GAGome was further used to calculate additional dependent features, including the total amount of CS and HS (as a sum of individually measured disaccharides), the CS and HS composition (as mass fractions of individual detectable CS and HS disaccharides) and the CS and HS charge (as the ratio of the sulfated disaccharides weighted by their charge and total CS and HS disaccharides).

### Statistical analysis

Before all analyses, GAGome features were transformed with Box-Cox transformation to identify and exclude outliers. Reference intervals were established for each urine and plasma free GAGome feature using a simple nonparametric method after outlier identification and exclusion. The lower and upper reference limits (reference intervals) for individual GAGs were estimated as the 2.5th and 97.5th percentiles of the distribution of measured values for the reference population, respectively.

The correlation between each GAGome feature and each clinical (e.g. age) or biochemical (e.g. LDL) variable was calculated by univariable linear regression of a given GAGome feature as response variable on a given clinical or biochemical variable as explanatory variable. We computed the statistical significance of each correlation using a likelihood ratio test versus an intercept-only regression model. Multiple hypothesis testing was adjusted using the Benjamini-Hochberg or Bonferroni corrections depending on if clinical or biochemical variables were tested, and *FDR* values < 0.1 and *p*-values < 1.67·10^-4^ were considered statistically significant, respectively. All statistical analyses were carried out in *R* (4.0.4) (*17*).

## Results

### Subject characteristics

We prospectively enrolled two cohorts of self-rated healthy adults with no history nor family history of cancer (except non-melanoma skin cancer) from one site in Stockholm, Sweden (Cohort 1, *N* = 292 and 2, *N* = 16), for a total of 308 participants (Table 1). Cohort 1 and 2 formed the reference sample group to establish reference intervals.

**Table 1.** Subject characteristics in the reference sample group (Cohort 1 and Cohort 2). Distributions are summarized as median and min-max range in brackets. Missing values were omitted.

|  | **Cohort 1** | **Cohort 2** | **Cohort 1 + 2** |
| --- | --- | --- | --- |
| **N** | 292 | 16 | 308 |
| **Age** | 57 (22-78) | 43 (27-51) | 57 (22-78) |
| **Gender** |  |  |  |
| Female | 183 | 5 | 188 |
| Male | 109 | 11 | 120 |
| **Self-rated health** |  |  |  |
| Moderate | 10 | 0 | 10 |
| Good | 153 | 5 | 158 |
| Very good | 129 | 11 | 140 |
| **Blood chemistry biomarkers** |  |  |  |
| ALAT ($\mu\text{kat/L}$) | 0.36 (0.13-2.44) | 0.49 (0.27-1.1) | 0.37 (0.13-2.44) |
| ASAT ($\mu\text{kat/L}$) | 0.41 (0.22-1.65) | 0.41 (0.27-0.59) | 0.41 (0.22-1.65) |
| Calcium ($\text{mmol/L}$) | 2.39 (2.15-2.72) | 2.42 (2.17-2.58) | 2.39 (2.15-2.72) |
| Creatinine ($\mu\text{mol/L}$) | 69 (46-167) | 80 (59-102) | 69.5 (46-167) |
| CRP ($\text{mg/L}$) | 1.4 (0.2-54.1) | 0.65 (0.27-5.5) | 1.3 (0.2-54.1) |
| EGFR ($\text{mL/min/1.73m}^2$) | 82.5 (33-90) | 90 (76-90) | 83 (33-90) |
| Glycated hemoglobin ($\text{mmol/mol}$) | - | 33.5 (28-39) | 33.5 (28-39) |
| HDL ($\text{mg/dL}$) | 61.87 (29.78-123.74) | - | 61.87 (29.78-123.74) |
| LDL ($\text{mg/dL}$) | 131.48 (46.4-239.75) | - | 131.48 (46.4-239.75) |
| Potassium ($\text{mmol/L}$) | 4.2 (3.5-5.2) | 4.2 (3.9-4.9) | 4.2 (3.5-5.2) |
| Prostate specific antigen level ($\text{ng/mL}$) | 0.8 (0.11-9.8) | 0.7 (0.33-1.4) | 0.8 (0.11-9.8) |
| Sodium ($\text{mmol/L}$) | 39 (132-145) | 140.5 (138-144) | 139 (132-145) |
| **Complete blood count** |  |  |  |
| Hematocrit (%) | 0.42 (0.35-0.51) | 0.44 (0.37-0.48) | 0.42 (0.35-0.51) |
| Hemoglobin ($\text{g/L}$) | 141 (116-178) | 145.5 (118-163) | 141 (116-178) |
| Mean corpuscular hemoglobin ($\text{pg}$) | 30 (24-37) | 30 (27-31) | 30 (24-37) |
| Mean corpuscular hemoglobin concentration ($\text{g/L}$) | 336 (304-366) | 331 (315-351) | 336 (304-366) |
| Mean corpuscular volume ($\text{fL}$) | 90 (81-101) | 89.5 (86-94) | 90 (81-101) |
| Absolute neutrophil count ($10^9/\text{L}$) | 3.2 (1-7.7) | - | 3.2 (1-7.7) |
| Basophil count ($10^9/\text{L}$) | 0 (0-0.2) | - | 0 (0-0.2) |
| Eosinophil count ($10^9/L$) | 0.1 (0-2.8) | - | 0.1 (0-2.8) |
| Erythrocyte count ($10^{12}/L$) | 4.6 (3.8-5.8) | 4.85 (4.1-5.3) | 4.6 (3.8-5.8) |
| Leukocyte count ($10^9/L$) | 5.6 (2.8-10.8) | 5.75 (3.5-7.7) | 5.6 (2.8-10.8) |
| Lymphocyte count ($10^9/L$) | 1.7 (0.7-4.5) | - | 1.7 (0.7-4.5) |
| Monocyte count ($10^9/L$) | 0.4 (0.2-0.9) | - | 0.4 (0.2-0.9) |
| Platelet count ($10^9/L$) | 261 (137-488) | 239 (166-377) | 261 (137-488) |
| **Blood chemistry status** |  |  |  |
| No abnormal values | 135 | 11 | 146 |
| 1-2 abnormal values | 134 | 5 | 139 |
| >2 abnormal values | 23 | 0 | 23 |

Subjects’ characteristics were balanced between cohorts (Table 1). Across cohorts, the median age was 57 years (range: 22 - 78) with 188 (61%) women and 120 (39%) men. Virtually all subjects self-reported good (51%) or very good (45%) health status. The panel of blood chemistry markers measured in Cohort 1 and 2 was the same, except for HDL/LDL (measured in Cohort 1 only) and glycated hemoglobin (measured in Cohort 2 only). Complete blood counts were available for Cohort 1, while Cohort 2 had available measurements for erythrocyte, leukocyte and platelet counts. Blood chemistry showed normal values for all biomarkers in 47% of the subjects, values outside the reference intervals for up to 2 biomarkers in 45% of the subjects, and more than two abnormal values in 23 subjects (8%). The abnormal blood chemistry values were mostly due to elevated calcium (12.2 % subjects), low sodium (10.4%), low CRP (7.5%), low EGFR (7.2%) and elevated PSA (5%). We kept subjects with abnormal values in the reference sample group and performed a sensitivity analysis of the reference values within this subgroup.

The plasma and urine free GAGomes were measured in a single GLP-compliant blinded laboratory using a standardized kit on UHPLC-MS/MS in all Cohort 1 and 2 samples (*N* = 308).

### Effect of blood chemistry on the normal free GAGome

We sought to characterize whether other blood chemistry biomarkers were correlated with the normal urine or plasma free GAGome.

First, we classified subjects in groups indicative of their general health status depending on the number of abnormal values for the blood chemistry biomarkers here tested. Specifically, we grouped the subjects into three groups: 1) no abnormal values (“No abnormal value”, *N* = 146), 2) one or two abnormal values (“1-2 abnormal values”, *N* = 139), or 3) more than two abnormal values (“>2 abnormal values”, *N* = 23). We did not observe any statistical associations between any of GAGome features with any of these groups (Table S1).

Second, we investigated linear correlations between the concentration of each detectable GAGome feature and each of the 25 blood chemistry biomarker level (as a continuous variable) and focused on correlations that reached statistical significance after controlling for multiple testing (*p* < 1.67·10^-4^, Bonferroni correction, Figure S2). In urine, we observed weak to moderate positive correlations (ρ = 0.23-0.3) between multiple GAGome features with creatinine. Additionally, the urine 6s CS concentration was positively correlated with hemoglobin, hematocrit, and erythrocyte count (ρ = 0.24 - 0.25) and negatively correlated with HDL (ρ = −0.22). For the plasma free GAGome, the total plasma and 4s CS concentrations were weakly positively correlated with hemoglobin and erythrocyte count (ρ = 0.21-0.28). Additionally, plasma 4s CS was correlated with hematocrit (ρ = 0.25), while 0s CS negatively correlated with HDL at ρ = −0.22, (Figure S2).

### Effect of age and gender on the normal free GAGome

We sought to identify if the urine or plasma free GAGome correlated with age and gender that could partition the reference intervals.

We did not observe any statistically significant association between age (as a continuous variable) and any GAGome feature (Table S2, Figure S1).

We observed statistically significant associations between gender and 10 detectable GAGome features (FDR < 0.1, Table S3, Figure 1-2). In plasma, the effect of gender was generally limited to an average 7% increase in total CS and 11% increase 4s CS concentration in males. The urine of males contained on average 31%-41% higher concentration for the major CS disaccharides (0s, 4s, 6s, and 2s6s CS), resulting in an average 34% increase in total CS; and an average 50% and 73% higher concentration for the major HS disaccharides (0s and Ns, respectively), resulting in an average 47% increase in total HS.

**Figure 1.**
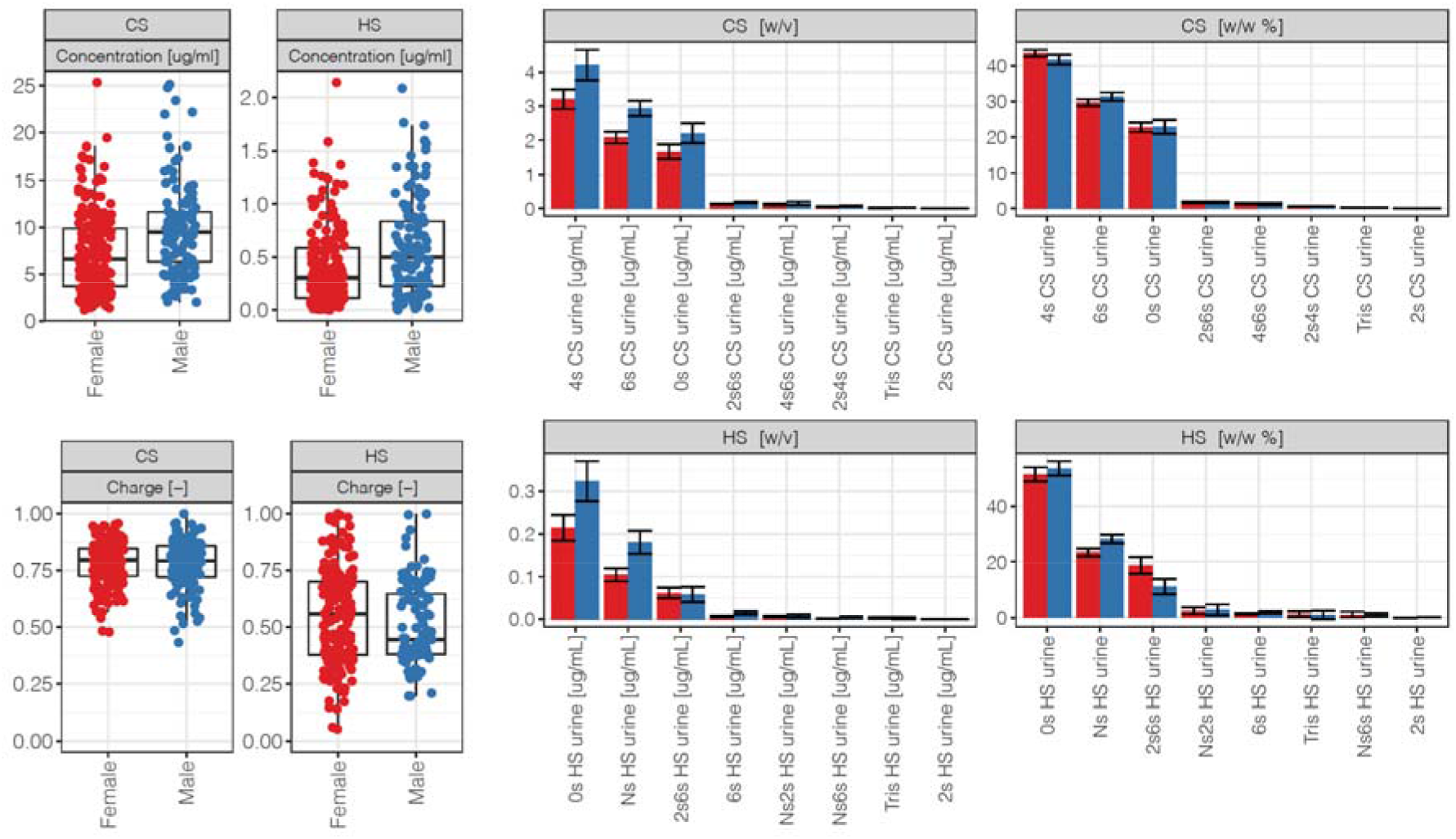
Urine free total CS and HS concentration (μg/ml), charge, and disaccharide concentration (μg/ml) and composition (in mass fraction %) in the reference sample group by gender (Cohort 1 and 2, *N_females_* = 188 *N_males_* = 120). Error bars indicate +/− 1.96 SEM (95% confidence interval). Key: Red – female, blue - male.

**Figure 2.**
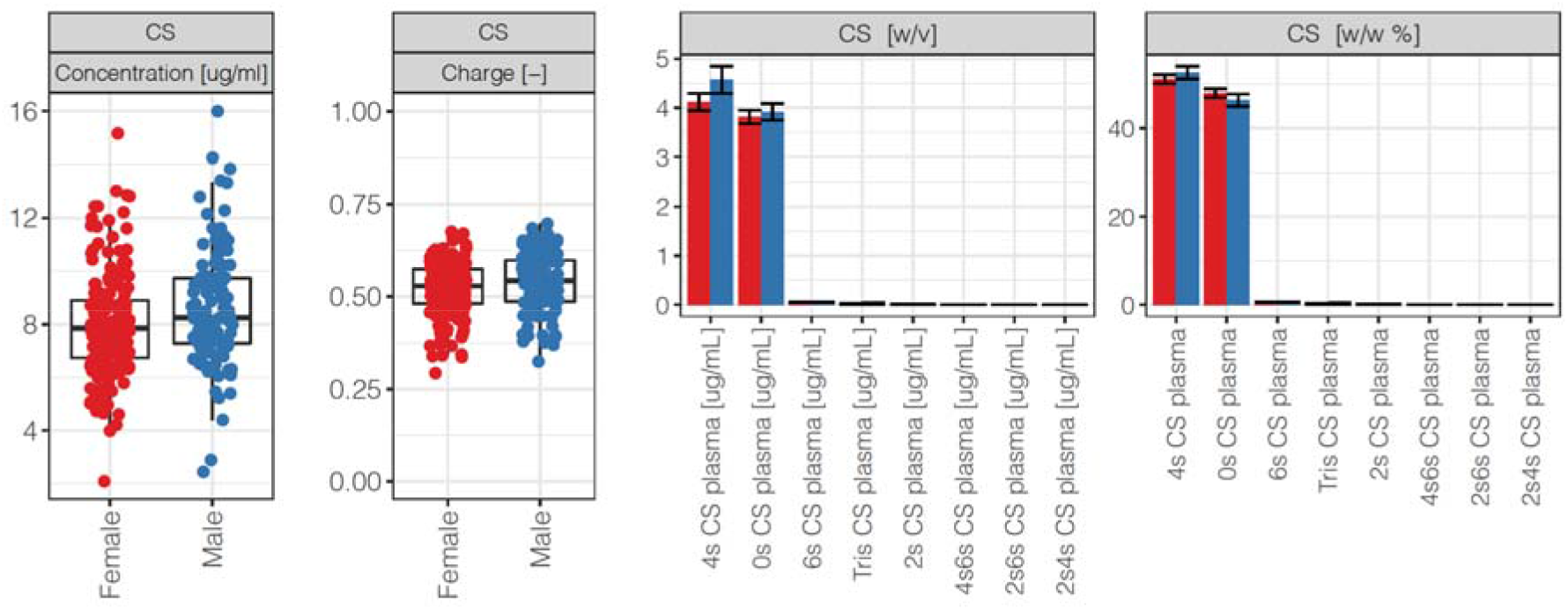
Plasma free total CS concentration (μg/ml), charge, and disaccharide concentration (μg/ml) and composition (in mass fraction %) in the reference sample group (Cohort 1 and 2). Plasma free HS was undetectable. Error bars indicate +/- 1.96 SEM (95% confidence interval). Key: Red – female, blue - male.

### Reference intervals for the normal free GAGome

Given the effects of gender on the GAGome, we decided to partition reference intervals by gender. We thus defined reference intervals for urine and plasma free GAGome concentrations and composition for apparently healthy males and females between the ages of 22 and 78.

First, we established reference intervals for the urine free GAGome after outlier identification and exclusion. For each disaccharide, *bona fide* outliers were identified according to a pre-specified procedure and omitted (median % of outliers across CS disaccharides: 2% in males, 0% in females; across HS: 3% in males, 3% in females).

We then established the reference intervals partitioned by gender of all urine free CS and HS features (Figure 1, Table 2).

**Table 2.**
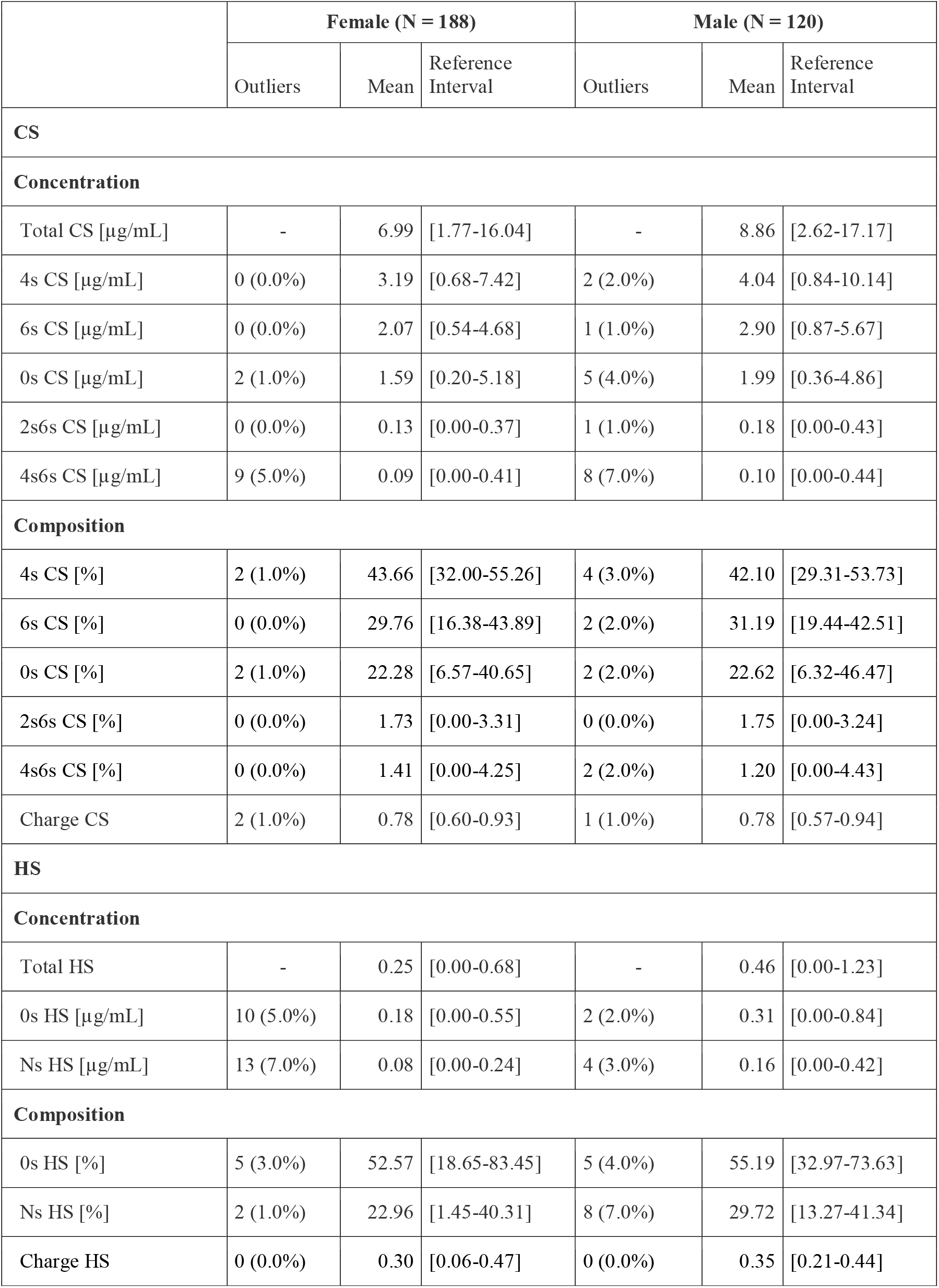
Reference intervals of urine free total CS and HS concentration (μg/ml) and disaccharide concentration (μg/ml) and composition (% w/w) by gender. Outliers were excluded.

|  | Female (N = 188) |  |  | Male (N = 120) |  |  |
| --- | --- | --- | --- | --- | --- | --- |
|  | Outliers | Mean | Reference Interval | Outliers | Mean | Reference Interval |
| **CS** |  |  |  |  |  |  |
| **Concentration** |  |  |  |  |  |  |
| Total CS [µg/mL] | - | 6.99 | [1.77-16.04] | - | 8.86 | [2.62-17.17] |
| 4s CS [µg/mL] | 0 (0.0%) | 3.19 | [0.68-7.42] | 2 (2.0%) | 4.04 | [0.84-10.14] |
| 6s CS [µg/mL] | 0 (0.0%) | 2.07 | [0.54-4.68] | 1 (1.0%) | 2.90 | [0.87-5.67] |
| 0s CS [µg/mL] | 2 (1.0%) | 1.59 | [0.20-5.18] | 5 (4.0%) | 1.99 | [0.36-4.86] |
| 2s6s CS [µg/mL] | 0 (0.0%) | 0.13 | [0.00-0.37] | 1 (1.0%) | 0.18 | [0.00-0.43] |
| 4s6s CS [µg/mL] | 9 (5.0%) | 0.09 | [0.00-0.41] | 8 (7.0%) | 0.10 | [0.00-0.44] |
| **Composition** |  |  |  |  |  |  |
| 4s CS [%] | 2 (1.0%) | 43.66 | [32.00-55.26] | 4 (3.0%) | 42.10 | [29.31-53.73] |
| 6s CS [%] | 0 (0.0%) | 29.76 | [16.38-43.89] | 2 (2.0%) | 31.19 | [19.44-42.51] |
| 0s CS [%] | 2 (1.0%) | 22.28 | [6.57-40.65] | 2 (2.0%) | 22.62 | [6.32-46.47] |
| 2s6s CS [%] | 0 (0.0%) | 1.73 | [0.00-3.31] | 0 (0.0%) | 1.75 | [0.00-3.24] |
| 4s6s CS [%] | 0 (0.0%) | 1.41 | [0.00-4.25] | 2 (2.0%) | 1.20 | [0.00-4.43] |
| Charge CS | 2 (1.0%) | 0.78 | [0.60-0.93] | 1 (1.0%) | 0.78 | [0.57-0.94] |
| **HS** |  |  |  |  |  |  |
| **Concentration** |  |  |  |  |  |  |
| Total HS | - | 0.25 | [0.00-0.68] | - | 0.46 | [0.00-1.23] |
| 0s HS [µg/mL] | 10 (5.0%) | 0.18 | [0.00-0.55] | 2 (2.0%) | 0.31 | [0.00-0.84] |
| Ns HS [µg/mL] | 13 (7.0%) | 0.08 | [0.00-0.24] | 4 (3.0%) | 0.16 | [0.00-0.42] |
| **Composition** |  |  |  |  |  |  |
| 0s HS [%] | 5 (3.0%) | 52.57 | [18.65-83.45] | 5 (4.0%) | 55.19 | [32.97-73.63] |
| Ns HS [%] | 2 (1.0%) | 22.96 | [1.45-40.31] | 8 (7.0%) | 29.72 | [13.27-41.34] |
| Charge HS | 0 (0.0%) | 0.30 | [0.06-0.47] | 0 (0.0%) | 0.35 | [0.21-0.44] |

The average urine free total CS concentration was 8.86 μg/ml in males and 6.99 μg/ml in females. The CS composition in males and females was nearly identical. The three major disaccharides made up 42% and 44% (4s CS), 31% and 30% (6s CS), and 23% and 22% (0s CS) of the CS fraction in urine of males and females, respectively. Of the multi-sulfated CS disaccharides, the 2s6s CS and 4s6s CS cumulatively contributed approximately 3% to the CS fraction in both males and females, while 2s4s CS and Tris CS were undetectable. The average urine CS charge was 0.78 for males and females.

The average urine total free HS concentration was 0.46 μg/ml in males and 0.25 μg/ml in females. The only detectable free HS disaccharides were 0s HS and Ns HS. The HS urine composition was 55% 0s HS and 30% Ns HS in males, and 53% 0s HS and 23% Ns HS in females. The average HS charge was 0.35 in males and 0.30 in females.

We repeated the procedure above for the plasma free GAGome (median % of outliers across CS disaccharides: 0 % in males or 1% in females, Figure 2, Table 3). The plasma HS fraction was largely undetectable (mean total HS < 0.001 μg/ml) and we therefore omitted it from further analyses.

**Table 3.** Reference intervals of plasma free total CS concentration (μg/ml) and disaccharide concentration (μg/ml) and composition (% w/w) by gender. Plasma free HS was undetectable. Outliers were excluded.

|  | Female (N = 188) |  |  | Male (N = 120) |  |  |
| --- | --- | --- | --- | --- | --- | --- |
|  | Outliers | Mean | Reference Interval | Outliers | Mean | Reference Interval |
| **CS** |  |  |  |  |  |  |
| **Concentration** |  |  |  |  |  |  |
| Total CS |  | 7.79 | [4.91-11.28] |  | 8.52 | [5.93-12.78] |
| 4s CS [µg/mL] | 6 (3.0%) | 4.07 | [2.07-6.37] | 2 (2.0%) | 4.55 | [2.25-8.02] |
| 0s CS [µg/mL] | 4 (2.0%) | 3.78 | [2.26-5.48] | 4 (3.0%) | 3.96 | [2.56-5.82] |
| **Composition** |  |  |  |  |  |  |
| 4s CS [%] | 2 (1.0%) | 51.41 | [36.16-62.06] | 0 (0.0%) | 52.64 | [36.64-65.46] |
| 0s CS [%] | 1 (1.0%) | 48.02 | [37.77-64.12] | 0 (0.0%) | 46.37 | [33.92-62.09] |
|  | Outliers | Mean | Reference Interval | Outliers | Mean | Reference Interval |
| Charge CS | 0 (0.0%) | 0.52 | [0.36-0.62] | 0 (0.0%) | 0.53 | [0.37-0.66] |

The average plasma free total CS concentration was 8.52 μg/ml in males and 7.79 μg/ml in females. The CS composition was nearly identical across genders, 53% and 51% 4s CS and 46% and 48% 0s CS (Table 3) for males and females, respectively. The remaining free CS disaccharides were undetectable. The average plasma CS charge was 0.53 in men and 0.52 in women.

### Transference of reference intervals in an independent population

We validated the transference of the above established reference intervals for each GAGome feature in two independent populations from two distinct geographical sites (Cohort 3 and 4) by determining the complete urine and plasma free GAGome in Cohort 3 (N = 110, 60 males and 50 females) and Cohort 4 (N = 30, 15 males and 15 females). The average age was 59 years old (range 30-84) for Cohort 3 and 45 years old (range 22-66) for Cohort 4. We observed that Cohort 3 and 4 had largely similar GAGome measurements. Therefore, we opted to carry out the transference analysis in a group combining Cohort 3 and Cohort 4, partitioned by gender (*N* = 140, 65 females and 75 males, see Tables S4 and S5 for transference analyses on separate cohorts).

Across all urine free GAGome features, we excluded 0-9.2 % (median = 1.5%) outliers in females and 0-12 % (median = 3.3%) in males. Across all plasma free GAGome features, we excluded 0-4.6 % (median = 1.5%) outliers in females and 0-1.3 % (median = 0%) in males. We next determined the percentage of values outside the established reference limits for each urine and plasma GAGome feature, where <5% was considered acceptable for transference validation.

In urine, we observed that the transference of reference intervals was validated in both genders for the concentration of all detectable free GAGome features (Table 4). The total urine CS was outside the reference interval in 5.6% samples, while all samples had total HS within the reference interval. As regards composition, we observed a shift towards a higher 4s CS % (mean in reference sample group: 44% in females, 42% in males; in transference group: 50% in females, 47% in males) and concomitantly other smaller shifts towards lower values in the remaining CS disaccharides.

**Table 4.**
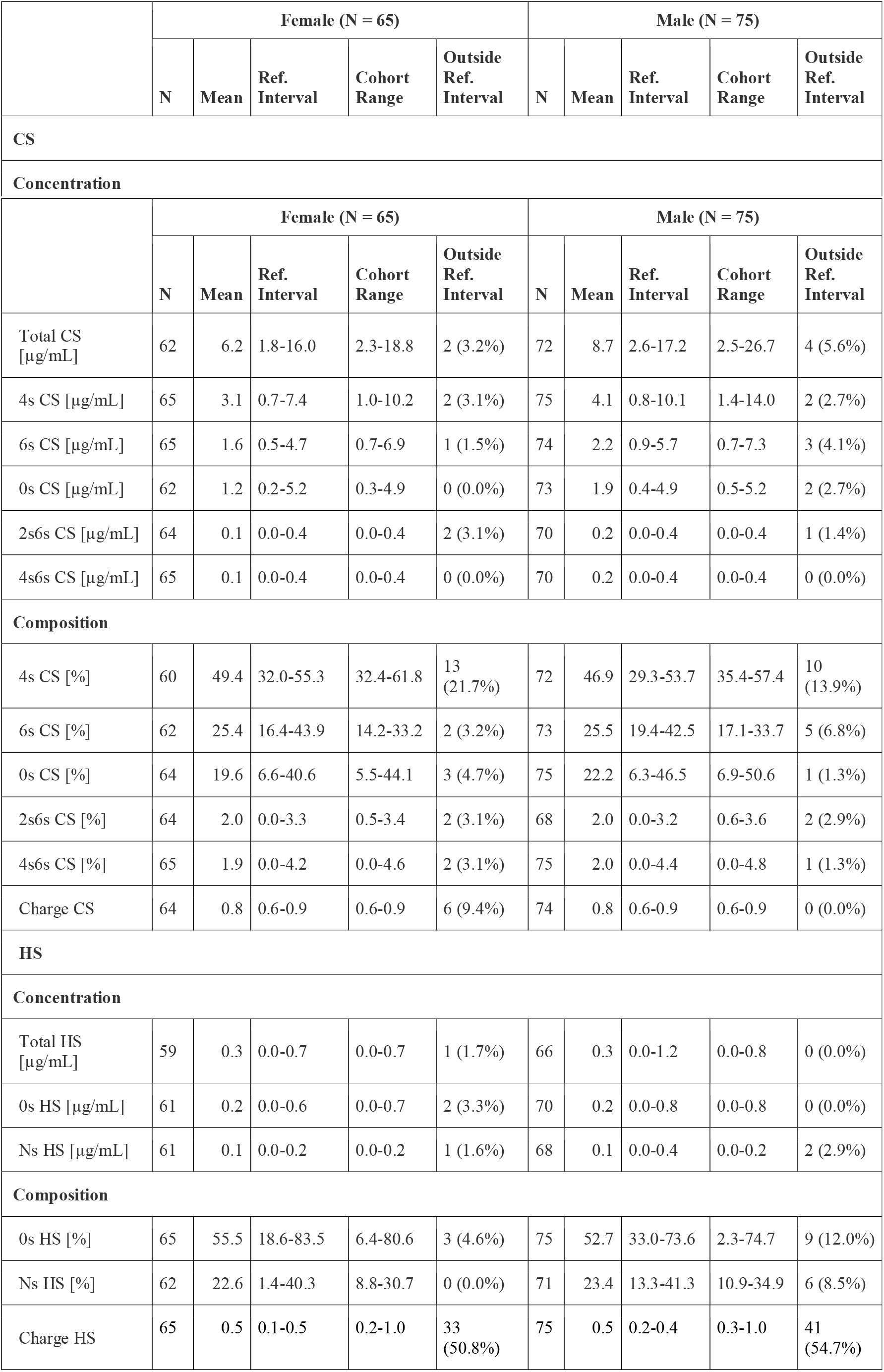
Transference of reference intervals of urine free CS and HS in an independent population (Cohorts 3 and 4). Outliers were excluded.

In plasma, we could not validate the transference of reference intervals for either of the two detectable plasma free GAGome features (0s CS and 4s CS) since 5 to 29% of the transference group had increased concentration (Table 5). The discrepancy was less pronounced for composition, where 11-22% had values outside the reference intervals.

**Table 5.** Transference of reference intervals of plasma free CS in an independent population (Cohorts 3 and 4). Outliers were excluded.

|  | Female (N = 65) |  |  |  |  | Male (N = 75) |  |  |  |  |
| --- | --- | --- | --- | --- | --- | --- | --- | --- | --- | --- |
|  | N | Mean | Ref. Interval | Cohort Range | Outside Ref. Interval | N | Mean | Ref. Interval | Cohort Range | Outside Ref. Interval |
| **CS** |  |  |  |  |  |  |  |  |  |  |
| **Concentration** |  |  |  |  |  |  |  |  |  |  |
| Total CS [µg/mL] | 62 | 10.0 | 4.9-11.3 | 6.7-14.9 | 18 (29.0%) | 74 | 10.1 | 5.9-12.8 | 6.7-15.9 | 4 (5.4%) |
| 4s CS [µg/mL] | 63 | 5.6 | 2.1-6.4 | 3.2-8.5 | 18 (28.6%) | 75 | 5.5 | 2.3-8.0 | 2.9-11.0 | 4 (5.3%) |
| 0s CS [µg/mL] | 64 | 4.4 | 2.3-5.5 | 2.6-8.5 | 7 (10.9%) | 74 | 4.5 | 2.6-5.8 | 2.4-8.2 | 12 (16.2%) |
| **Composition** |  |  |  |  |  |  |  |  |  |  |
| 4s CS [%] | 65 | 56.3 | 36.2-62.1 | 30.0-72.3 | 14 (21.5%) | 75 | 54.6 | 36.6-65.5 | 33.7-72.8 | 11 (14.7%) |
| 0s CS [%] | 64 | 42.8 | 37.8-64.1 | 26.9-59.1 | 14 (21.9%) | 75 | 44.7 | 33.9-62.1 | 26.4-65.9 | 11 (14.7%) |
| Charge CS | 65 | 0.6 | 0.4-0.6 | 0.3-0.7 | 15 (23.1%) | 75 | 0.6 | 0.4-0.7 | 0.3-0.7 | 12 (16.0%) |

## Discussion

In this study, we established reference intervals for the urine and plasma free GAGomes in a large adult healthy population by taking advantage of standardized high throughput UHPLC-MS/MS method (*15*). In addition, the extensive demographic and biochemical characterization of subjects allowed us to assess novel correlations with the free GAGome.

We found that no free GAGome feature showed any notables differences with respect to age in adults. While this finding in urine agree with previous studies analyzing total – as opposed to free - GAGomes (*18*), it contradicts similar studies in plasma, where an increase with age was reported in males only (*19*). In contrast to age, we found that the concentrations of several CS and HS disaccharides were higher in males than females with a larger effect in urine than plasma. Previous studies noted gender differences in total GAGomes, although in the opposite directions for both urine and plasma (*18*, *19*). The origin of these discrepancies remains to be ascertained, but we speculate that they could be attributed to the focus on the total rather than the free GAGome fraction in previous studies, as well as older analytical techniques for GAG measurements and smaller sample sizes. We also observed that urine free GAGomes appeared independent of other blood chemistry biomarkers, including markers of inflammation, glucose metabolism, and liver functions – underscoring that they may reflect an independent physiological state in adults. The only exception was serum creatinine, which in the context of its correlation with urine free GAGomes may indicate a dependency on urine excretion rather than renal function. Thus, we speculate that normalization by urinary creatinine could render urine free GAGome measurements more robust. In contrast, plasma free CS had weak correlations with markers related to erythrocytes, particularly 4s CS. This association has been previously described for platelets where their activation can lead to rapid increases of circulating 4s CS (*20*), but not in the context of other blood cells such as erythrocytes.

We observed that the reference intervals established for free GAGomes were remarkably tight in the reference sample group. In plasma, each free GAGome feature deviated by ~75% at most from the mean. Similarly, in the urine each feature deviated by a maximum of ~3-fold from the mean. These findings suggest that urine and plasma free GAGomes have stable and predictable levels in a healthy adult. This conclusion appears corroborated by the transference analysis that validated all reference intervals established for urine in independent adult samples from two different geographical sites. Even among the free GAGome features that failed to validate the reference intervals, chiefly in plasma, the deviation from the reference sample group was limited. For example, the mean plasma 4s CS concentration was 4.1 ng mL^-1^ in the female reference sample group and 5.6 ng mL in the transference group, that is 36% higher. In comparison, the reference interval for the platelet count in the reference sample group spans 137 to 488•10^9^ L^-1^, a 3-fold difference between the reference limits. Considering this, we speculation that if a substantial deviation of a free GAGome feature from the here reported reference interval is observed in an adult, then this deviation may be more indicative of a disease state than physiological variability. This makes free GAGomes suitable candidates for biomarker studies.

In conclusion, this study established and validated reference intervals for plasma and urine free GAGomes in the largest adult population to date. As such, we believe that this study represents a critical resource for physiology and biomarker research using biofluidic free GAGomes.

## Acknowledgments

This work was supported by the Knut and Alice Wallenberg Foundation, Cancerfonden, and the IngaBritt och Arne Lundbergs Forskningsstiftelse (J.N.) and in part by the European Union’s Horizon 2020 research and innovation programme under grant agreement No 849251 and the EIT Healthy 2019 Digital Sandbox (Grant No 2019-DS1001-6543) (Fr.G.). The Lifelines initiative has been made possible by the subsidy from the Dutch ministry of Health, Welfare, and Sport, the Dutch Ministry of Economic Affairs, the University Medical Center Groningen (UMCG), Groneningen University, and the Provinces in the North of the Netherlands (Drenthe, Friesland, Groningen). The authors wish to thank the research nurses Lena Solitander and Elisabeth Kapocs, and Karolinska Trial Alliance for help in the coordination, supervision, and collection of clinical data and samples used in this study.

## Author contributions

Fr.G., J.N conceived, designed, and coordinated the study. S.B., A.L., and Fr.G. performed data analysis and statistics. N.V., Fa.G., and F.M. assisted with the laboratory method. M.L. contributed to subject enrollment. S.B., A.L., and Fr.G. drafted the manuscript. All authors edited and approved the manuscript in its final form.

