## Supplementary information for "Analysis of Normal Levels of Urine and Plasma Free Glycosaminoglycans in Adults"

**Affiliations:**
[1] Department of Biology and Biological Engineering, Chalmers University of Technology, 412 96 Göteborg, Sweden.
[2] Department of Life Sciences, University of Modena and Reggio Emilia, 411 25 Modena, Italy.
[3] Department of Molecular and Clinical Medicine/Wallenberg Laboratory, Institute of Medicine, Sahlgrenska Academy, University of Gothenburg, Gothenburg, Sweden.
[4] Departments of Oncology, Sahlgrenska Academy, University of Gothenburg, 41345 Göteborg, Sweden.
[5] Department of Oncology, Sahlgrenska University Hospital, 413 45, Gothenburg, Sweden.
[6] BioInnovation Institute, Ole Maaløes Vej 3, DK 2200 Copenhagen N, Denmark.

[*] Current affiliation: Elypta AB, 171 65 Stockholm, Sweden.

**Contents:**
Figure S1 -S2
Tables S1- S5

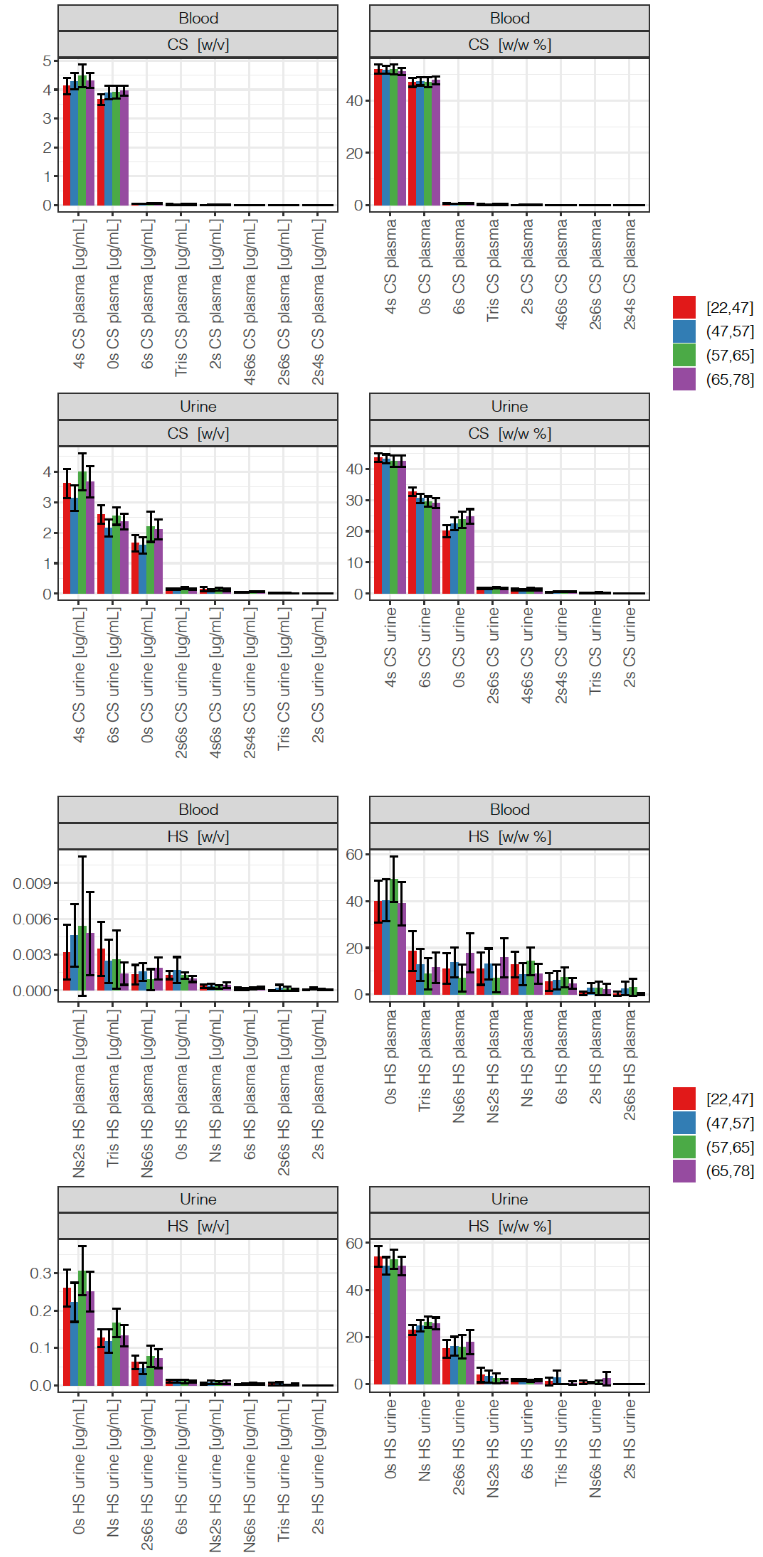


**Figure S1.** CS (top) and HS (bottom) disaccharide concentration (µg/ml) and composition (mass fraction %) across age groups (Cohort 1 and 2, $N = 308$). Error bars indicate +/- 1.96 SEM (95% confidence interval).

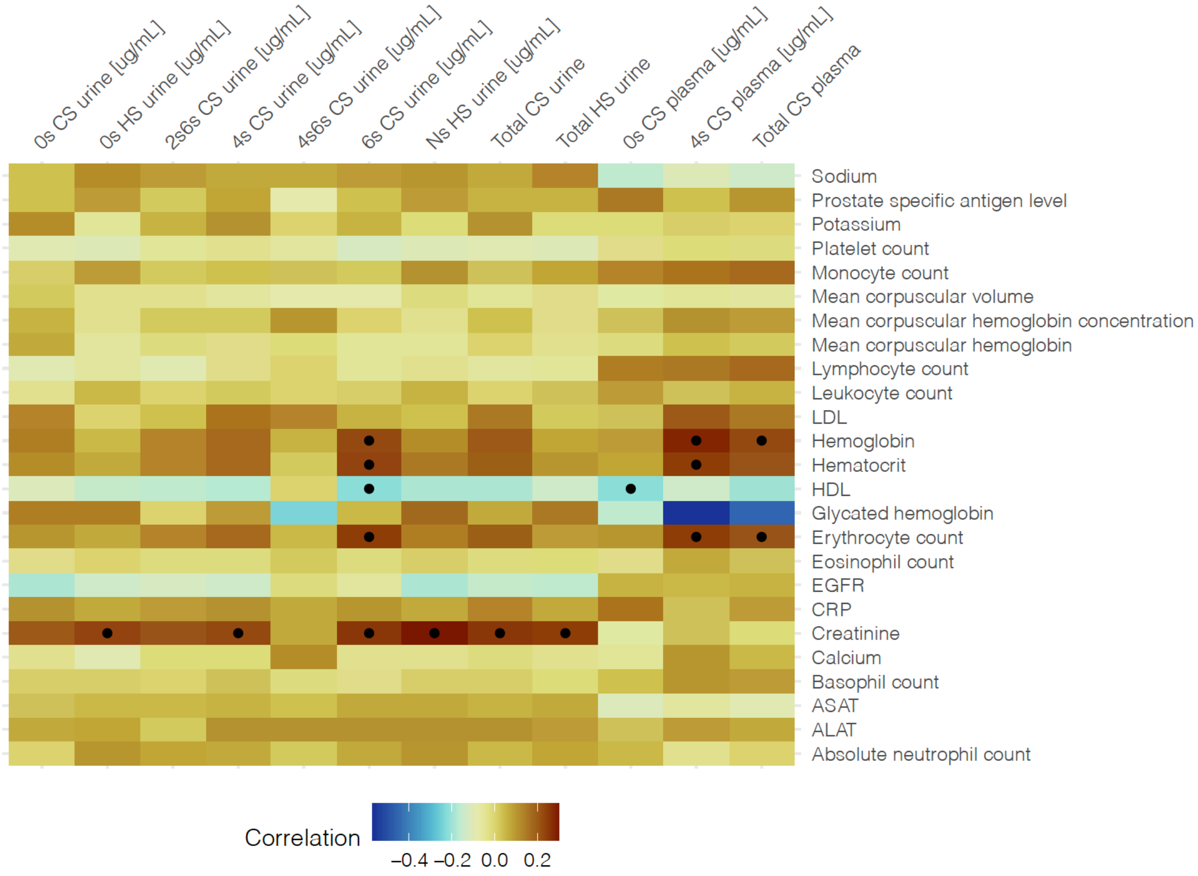


**Figure S2.** Correlation matrix (in terms of Pearson correlation coefficient) between detectable glycosaminoglycan disaccharides and total CS/HS (µg/ml) and 25 blood biomarkers (Cohort 1 and 2, $N$ = 308 – note that LDL/HDL was measured only in Cohort 1, while glycated hemoglobin was measured only in Cohort 2). Black dots indicate significant correlations after adjusting for multiple testing ($p < 1.67 \cdot 10^{-4}$, Bonferroni correction).

**Table S1.** Change in the total concentration (µg/ml) and disaccharide concentration (µg/ml) for CS and HS across groups with different counts of abnormal blood chemistry biomarker levels. FDR values < 0.1 were considered significant.

| | **Change if 1-2 abnormal lab values (%)** | **Change if >2 abnormal lab values (%)** | **FDR** |
|---|---|---|---|
| **Plasma CS** | | | |
| Total CS plasma [µg/mL] | 4 | 5 | 0.27 |
| 0s CS plasma [µg/mL] | 5 | 3 | 0.44 |
| 4s CS plasma [µg/mL] | 4 | 7 | 0.26 |
| **Urine HS** | | | |
| Total HS urine | 29 | 13 | 0.17 |
| Ns HS urine [µg/mL] | 39 | 19 | 0.14 |
| 0s HS urine [µg/mL] | 24 | 10 | 0.26 |
| **Urine CS** | | | |
| Total CS urine [µg/mL] | 20 | 9 | 0.17 |
| 0s CS urine [µg/mL] | 21 | 0 | 0.20 |
| 6s CS urine [µg/mL] | 16 | 11 | 0.17 |
| 4s CS urine [µg/mL] | 23 | 13 | 0.17 |
| 2s6s CS urine [µg/mL] | 23 | 14 | 0.17 |
| 4s6s CS urine [µg/mL] | 17 | −1 | 0.71 |

**Table S2.** Change in the total concentration (µg/ml) and disaccharide concentration (µg/ml) for CS, HS, and HA by year of age (Cohort 1 and 2, $N = 308$). FDR values < 0.1 were considered significant.

| | **Change per year (%)** | **FDR** |
|---|---:|---:|
| **Plasma CS** | | |
| Total CS plasma [µg/mL] | 0.2 | 0.438 |
| 0s CS plasma [µg/mL] | 0.2 | 0.438 |
| 4s CS plasma [µg/mL] | 0.2 | 0.505 |
| **Urine HS** | | |
| Total HS urine [µg/mL] | 0.5 | 0.505 |
| Ns HS urine [µg/mL] | 0.8 | 0.505 |
| 0s HS urine [µg/mL] | 0.4 | 0.563 |
| **Urine CS** | | |
| Total CS urine [µg/mL] | 0.3 | 0.505 |
| 0s CS urine [µg/mL] | 1.6 | 0.229 |
| 6s CS urine [µg/mL] | −0.2 | 0.505 |
| 4s CS urine [µg/mL] | 0.3 | 0.528 |
| 2s6s CS urine [µg/mL] | 0.3 | 0.563 |
| 4s6s CS urine [µg/mL] | 0.3 | 0.710 |

**Table S3.** Change in the total concentration (µg/ml) and disaccharide concentration (µg/ml) for CS, HS, and HA in male vs. females (Cohort 1 and 2, $N = 308$, 39% male). FDR values < 0.1 were considered significant.

| | **Change in Male (%)** | **FDR** |
|---|---:|---:|
| **Plasma CS** | | |
| Total CS plasma [µg/mL] | **7** | **0.018** |
| 0s CS plasma [µg/mL] | 3 | 0.337 |
| 4s CS plasma [µg/mL] | **11** | **0.007** |
| **Urine HS** | | |
| Total HS urine | **47** | **<0.001** |
| Ns HS urine [µg/mL] | **73** | **<0.001** |
| 0s HS urine [µg/mL] | **50** | **<0.001** |
| **Urine CS** | | |
| Total CS urine [µg/mL] | **34** | **<0.001** |
| 0s CS urine [µg/mL] | **32** | **0.005** |
| 6s CS urine [µg/mL] | **41** | **<0.001** |
| 4s CS urine [µg/mL] | **31** | **<0.001** |
| 2s6s CS urine [µg/mL] | **36** | **0.001** |
| 4s6s CS urine [µg/mL] | 35 | 0.133 |

**Table S4.** Transference of reference intervals of free urine and plasma CS and HS in an independent population (Cohort 3). Note that outliers were excluded.

| | Female (N=50) | | | | | Male (N=60) | | | | |
|---|---|---|---|---|---|---|---|---|---|---|
| | N | Mean | Range | Low | High | N | Mean | Range | Low | High |
| Urine Concentration | | | | | | | | | | |
| Total CS [µg/mL] | 49 | 5.34 | 2.3-11.4 | 0 (0.0%) | 0 (0.0%) | 59 | 7.80 | 3.6-17.5 | 0 (0.0%) | 1 (1.7%) |
| 4s CS [µg/mL] | 50 | 2.75 | 1.1-6.3 | 0 (0.0%) | 0 (0.0%) | 60 | 3.68 | 1.6-8.7 | 0 (0.0%) | 0 (0.0%) |
| 6s CS [µg/mL] | 50 | 1.30 | 0.7-2.4 | 0 (0.0%) | 0 (0.0%) | 59 | 1.95 | 0.9-3.8 | 0 (0.0%) | 0 (0.0%) |
| 0s CS [µg/mL] | 49 | 1.06 | 0.2-4.4 | 0 (0.0%) | 0 (0.0%) | 60 | 1.81 | 0.6-5.0 | 0 (0.0%) | 1 (1.7%) |
| 4s6s CS [µg/mL] | 49 | 0.12 | 0.0-0.4 | 0 (0.0%) | 0 (0.0%) | 57 | 0.17 | 0.0-0.4 | 0 (0.0%) | 0 (0.0%) |
| 2s6s CS [µg/mL] | 50 | 0.11 | 0.0-0.3 | 0 (0.0%) | 0 (0.0%) | 60 | 0.16 | 0.0-0.3 | 0 (0.0%) | 0 (0.0%) |
| Total HS [µg/mL] | 50 | 0.18 | 0.0-0.5 | 2 (4.0%) | 0 (0.0%) | 55 | 0.20 | 0.0-0.4 | 1 (1.8%) | 0 (0.0%) |
| 0s HS [µg/mL] | 50 | 0.13 | 0.0-0.4 | 1 (2.0%) | 0 (0.0%) | 60 | 0.16 | 0.0-0.4 | 1 (1.7%) | 0 (0.0%) |
| Ns HS [µg/mL] | 50 | 0.05 | 0.0-0.1 | 1 (2.0%) | 0 (0.0%) | 55 | 0.06 | 0.0-0.1 | 1 (1.8%) | 0 (0.0%) |
| Urine Composition | | | | | | | | | | |
| 4s CS | 49 | 50.56 | 38.4-61.8 | 0 (0.0%) | 13 (26.5%) | 59 | 46.53 | 35.4-57.4 | 0 (0.0%) | 7 (11.9%) |
| 6s CS | 50 | 24.77 | 14.2-32.7 | 2 (4.0%) | 0 (0.0%) | 59 | 24.84 | 17.1-33.5 | 5 (8.5%) | 0 (0.0%) |
| 0s CS | 50 | 18.42 | 5.5-37.9 | 1 (2.0%) | 0 (0.0%) | 59 | 21.96 | 9.7-39.4 | 0 (0.0%) | 0 (0.0%) |
| 4s6s CS | 48 | 2.23 | 0.9-4.4 | 0 (0.0%) | 1 (2.1%) | 59 | 2.22 | 0.5-4.3 | 0 (0.0%) | 0 (0.0%) |
| 2s6s CS | 49 | 2.03 | 1.1-3.4 | 0 (0.0%) | 2 (4.1%) | 54 | 1.93 | 1.1-2.9 | 0 (0.0%) | 0 (0.0%) |
| 0s HS | 50 | 55.18 | 6.4-74.8 | 3 (6.0%) | 0 (0.0%) | 60 | 53.69 | 2.3-74.7 | 5 (8.3%) | 2 (3.3%) |
| Ns HS | 47 | 22.41 | 12.2-30.7 | 0 (0.0%) | 0 (0.0%) | 59 | 23.25 | 10.9-34.9 | 6 (10.2%) | 0 (0.0%) |
| Plasma Concentration | | | | | | | | | | |
| Total CS [µg/mL] | 48 | 10.15 | 6.7-14.9 | 0 (0.0%) | 15 (31.2%) | 59 | 10.12 | 6.7-15.9 | 0 (0.0%) | 4 (6.8%) |
| 4s CS [µg/mL] | 49 | 5.62 | 3.2-8.5 | 0 (0.0%) | 14 (28.6%) | 60 | 5.35 | 2.9-11.0 | 0 (0.0%) | 2 (3.3%) |
| 0s CS [µg/mL] | 49 | 4.51 | 2.6-8.5 | 0 (0.0%) | 7 (14.3%) | 59 | 4.73 | 2.4-8.2 | 1 (1.7%) | 11 (18.6%) |
| Plasma Composition | | | | | | | | | | |
| 4s CS | 49 | 56.67 | 40.6-72.3 | 0 (0.0%) | 9 (18.4%) | 60 | 52.97 | 33.7-72.8 | 2 (3.3%) | 5 (8.3%) |
| 0s CS | 49 | 42.74 | 26.9-59.1 | 10 (20.4%) | 0 (0.0%) | 60 | 46.35 | 26.4-65.9 | 5 (8.3%) | 2 (3.3%) |

**Table S5.** Transference of reference intervals of free urine and plasma CS and HS in an independent population (Cohort 4). Note that outliers were excluded.

| | Male (N = 15) | | | | | Female (N = 15) | | | | |
|---|---|---|---|---|---|---|---|---|---|---|
| | N | Mean | Range | Low | High | N | Mean | Range | Low | High |
| Urine concentration | | | | | | | | | | |
| Total CS [µg/mL] | 13 | 13.73 | 5.2-27.3 | 0 (0.0%) | 3 (23.1%) | 14 | 8.95 | 3.6-19.4 | 0 (0.0%) | 2 (14.3%) |
| 4s CS [µg/mL] | 15 | 5.94 | 1.4-14.0 | 0 (0.0%) | 2 (13.3%) | 15 | 4.42 | 1.0-10.2 | 0 (0.0%) | 2 (13.3%) |
| 6s CS [µg/mL] | 15 | 3.18 | 0.7-7.3 | 2 (13.3%) | 1 (6.7%) | 15 | 2.59 | 1.0-6.9 | 0 (0.0%) | 1 (6.7%) |
| 0s CS [µg/mL] | 13 | 3.17 | 0.7-8.6 | 0 (0.0%) | 2 (15.4%) | 14 | 1.85 | 0.7-4.9 | 0 (0.0%) | 0 (0.0%) |
| 2s6s CS [µg/mL] | 15 | 0.26 | 0.0-0.7 | 0 (0.0%) | 3 (20.0%) | 15 | 0.21 | 0.0-0.6 | 0 (0.0%) | 3 (20.0%) |
| 4s6s CS [µg/mL] | 14 | 0.10 | 0.0-0.3 | 0 (0.0%) | 0 (0.0%) | 15 | 0.05 | 0.0-0.2 | 0 (0.0%) | 0 (0.0%) |
| Total HS [µg/mL] | 15 | 0.61 | 0.0-1.9 | 0 (0.0%) | 4 (26.7%) | 13 | 0.66 | 0.0-1.8 | 0 (0.0%) | 2 (15.4%) |
| 0s HS [µg/mL] | 15 | 0.41 | 0.0-1.3 | 0 (0.0%) | 4 (26.7%) | 13 | 0.43 | 0.0-1.3 | 0 (0.0%) | 2 (15.4%) |
| Ns HS [µg/mL] | 15 | 0.20 | 0.0-0.7 | 0 (0.0%) | 4 (26.7%) | 15 | 0.24 | 0.0-0.6 | 1 (6.7%) | 3 (20.0%) |
| Urine Composition | | | | | | | | | | |
| 4s CS | 15 | 48.59 | 29.7-66.6 | 0 (0.0%) | 4 (26.7%) | 15 | 42.04 | 24.1-63.6 | 3 (20.0%) | 1 (6.7%) |
| 6s CS | 14 | 28.01 | 21.8-33.7 | 0 (0.0%) | 0 (0.0%) | 14 | 29.21 | 22.1-35.7 | 0 (0.0%) | 0 (0.0%) |
| 0s CS | 15 | 21.40 | 6.9-42.9 | 0 (0.0%) | 0 (0.0%) | 15 | 27.00 | 8.3-69.7 | 0 (0.0%) | 3 (20.0%) |
| 2s6s CS | 15 | 1.96 | 0.1-3.7 | 0 (0.0%) | 1 (6.7%) | 15 | 2.11 | 0.5-3.9 | 0 (0.0%) | 1 (6.7%) |
| 4s6s CS | 14 | 0.68 | 0.0-2.4 | 0 (0.0%) | 0 (0.0%) | 14 | 0.48 | 0.0-2.7 | 0 (0.0%) | 0 (0.0%) |
| 0s HS | 15 | 56.39 | 29.5-80.6 | 0 (0.0%) | 0 (0.0%) | 15 | 48.70 | 16.5-73.7 | 1 (6.7%) | 1 (6.7%) |
| Ns HS | 14 | 24.41 | 19.4-30.5 | 0 (0.0%) | 0 (0.0%) | 12 | 24.34 | 15.8-29.4 | 0 (0.0%) | 0 (0.0%) |
| Plasma Concentration | | | | | | | | | | |
| Total CS [µg/mL] | 15 | 9.88 | 7.6-12.2 | 0 (0.0%) | 0 (0.0%) | 15 | 9.34 | 5.3-12.0 | 0 (0.0%) | 3 (20.0%) |
| 4s CS [µg/mL] | 15 | 6.15 | 4.1-8.5 | 0 (0.0%) | 2 (13.3%) | 15 | 5.41 | 2.3-7.6 | 0 (0.0%) | 4 (26.7%) |
| 0s CS [µg/mL] | 15 | 3.74 | 2.6-5.0 | 0 (0.0%) | 0 (0.0%) | 15 | 3.94 | 2.9-5.2 | 0 (0.0%) | 0 (0.0%) |
| Plasma Composition | | | | | | | | | | |
| 4s CS | 15 | 61.29 | 50.5-70.6 | 0 (0.0%) | 4 (26.7%) | 15 | 56.69 | 44.3-64.3 | 0 (0.0%) | 4 (26.7%) |
| 0s CS | 15 | 37.93 | 28.6-48.8 | 4 (26.7%) | 0 (0.0%) | 15 | 42.77 | 34.9-55.1 | 4 (26.7%) | 0 (0.0%) |
